# The synthetic histone-binding regulator protein PcTF activates interferon genes in breast cancer cells

**DOI:** 10.1101/186056

**Authors:** Kimberly C. Olney, David B. Nyer, Daniel A. Vargas, Melissa A. Wilson Sayres, Karmella A. Haynes

**Affiliations:** School of Biological and Health Systems Engineering, Arizona State University; School of Life Sciences, Arizona State University; The Biodesign Institute Center for Evolutionary Medicine, Arizona State University

**Keywords:** chromatin, breast cancer, Polycomb, tumor suppressor genes

## Abstract

Mounting evidence from genome-wide studies of cancer show that chromatin-mediated epigenetic silencing at large cohorts of genes is strongly linked to a poor prognosis. This mechanism is thought to prevent cell differentiation and enable evasion of the immune system. Drugging the cancer epigenome with small molecule inhibitors to release silenced genes from the repressed state has emerged as a powerful approach for cancer research and drug development. Targets of these inhibitors include chromatin-modifying enzymes that can acquire drug-resistant mutations. In order to directly target a generally conserved feature, elevated trimethyl-lysine 27 on histone H3 (H3K27me3), we developed the Polycomb-based Transcription Factor (PcTF), a fusion activator that targets methyl-histone marks via its N-terminal H3K27me3-binding motif, and co-regulates sets of silenced genes. Here, we report transcriptome profiling analyses of PcTF-treated breast cancer model cell lines. We identified a set of 19 PcTF-upregulated genes, or PUGs, that were consistent across three distinct breast cancer cell lines. These genes are associated with the interferon response pathway. Our results demonstrate for the first time a chromatin-mediated interferon-related transcriptional response driven by an engineered fusion protein that physically links repressive histone marks with active transcription.

## INTRODUCTION

In addition to DNA lesions, disruption of chromatin at non-mutated genes can support the progression of cancer. Chromatin is a dynamic network of interacting proteins, DNA, and RNA that organizes chromosomes within cell nuclei. These interactions regulate gene transcription and coordinate distinct, genome-wide expression profiles in different cell types. Chromatin mediates epigenetic inheritance [1,2] by regulating expression states that persist through cellular mitosis and across generations of sexually reproducing organisms [3,4]. Posttranslational modifications (PTMs) of histones within nucleosomes, the fundamental subunits of chromatin, play a central role in the epigenetic regulation of genes that control cell differentiation [5,6]. Several landmark studies have revealed that hyperactivity of the histone-methyltransferase enhancer of zeste 1 and 2 (EZH1, EZH2), which generates the histone PTM H3K27me3, is a feature shared by many types of cancer (recently reviewed in [7]). In breast cancer, elevated EZH2 has been linked to cell proliferation and metastasis [8,9] and a poor prognosis for breast cancer patients [10–13]. In stem cells and cancer cells, EZH2 generates H3K27me3 mark at nucleosomes (Fig. 1) near the promoters of developmental genes, represses transcription, and thus prevents differentiation to support the proliferative state in stem cells or neoplasia in cancer (reviewed in [5]). Polycomb Repressive Complex 1 (PRC1, also known as PRC1.2 or PRC1.4[14]) binds to the H3K27me3 mark through the polycomb chromodomain (PCD) motif of the CBX protein to stabilize the repressed state. Silencing is reinforced by other chromatin regulators including histone deacetylase (HDAC) and DNA methyltransferase (DMT) [15] (Fig. 1).

**Figure 1.**
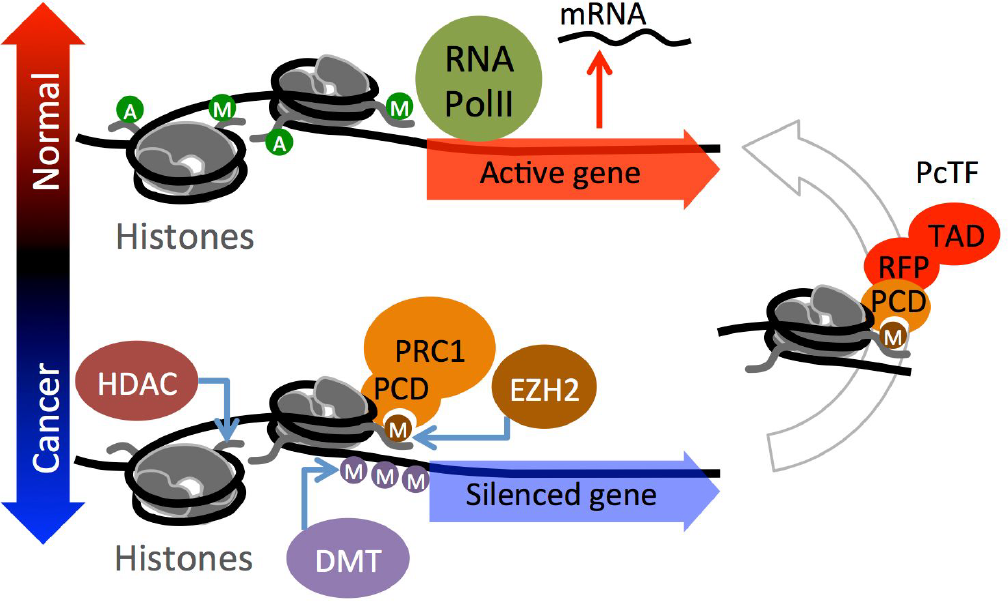
Reversal of a cancer-associated epigenetic state via the PcTF fusion protein. The lower half of the cartoon depicts the accumulation of repressive chromatin at a developmental gene. EZH2 generates H3K27me3, which is recognized by the PCD fold in the CBX protein of Polycomb Repressive Complex 1 (PRC1). Silencing is re-enforced by histone deacetylase (HDAC), and DNA methyltransferase (DMT) activity. The fusion protein PcTF contains an N-terminal PCD fold (cloned from CBX8) that binds H3K27me3 and stimulates transcription via its C-terminal activator domain to restore the active state (right side of the cartoon). A, acetylation; M, methylation; green circle, activation-associated PTM; orange or purple circle, repression-associated PTM; RFP, red fluorescent protein tag; TAD, transcriptional activation domain VP64.

The PRC module is a group of genes that is regulated by H3K27me3 and Polycomb transcriptional regulators [16,17]. Relatively high expression or upregulation of PRC module genes is associated with a non-proliferative state, cell adhesion, organ development, and normal anatomical structure morphogenesis [16]. Knockdown (depletion) of chromatin proteins (reviewed in [17,18]) and inhibition of Polycomb proteins with low molecular weight compounds [19–21] and peptides [22–24] stimulates expression of developmental genes and perturbs cancer-associated cell behavior. The interferon (IFN) pathway is often highly represented among silenced genes in cancer. IFN gene activity has been linked to apoptosis [25,26] and triggers the body’s immune system to attack cancer cells [27,28]. Decreased expression and increased levels of repressive epigenetic marks (e.g., DNA methylation) have been detected at IFN genes in Li–Fraumeni fibroblasts (39 of 85 silenced genes) [29], colon carcinomas [30], and triple negative breast cancers [31,32]. Transgenic overexpression of IFN1 in MCF7 breast cancer xenografts perturbs tumor growth in nude mice [26]. Treatment of cancerous cells with broad-acting epigenetic inhibitors of DNA methyltransferase (DNMTi) and histone deacetylase (HDACi) leads to activation of IFN genes which induces an arrest of cancer cell proliferation or sensitize cancer cells to immunotherapy [28,33,34].

The use of the FDA-approved DNA methyltransferase inhibitors (e.g., 5-azacytidine) to treat cancer, as well as the success of other epigenetic interventions in clinical trials [35,36] demonstrates that chromatin is a druggable target in cancer. Certain limitations of epigenetic inhibitor compounds could encumber complete efficacy of epigenetic therapy. Inhibitors do not interact directly with modified histones, indirectly activate silenced genes by blocking repressors, generate incomplete conversion of silenced chromatin into active chromatin [37,38], interact with off-target proteins outside of the nucleus [39], and do not affect resistant Polycomb protein mutants [40–42]. These limitations could be addressed by technologies that directly target H3K27me3 within the chromatin fiber. H3K27me3 is a highly conserved feature in cancers[7]. Even in cases where H3K27 becomes mutated to methionine in one allele[43,44], methylation of the wild-type copy of H3K27 is still present at repressed loci in cancer cells[45,46].

Our group developed a fusion protein called Polycomb-based Transcription Factor (PcTF), which specifically binds H3K27me3 [47] and recruits endogenous transcription factors to PRC-silenced genes (Fig. 1). In bone, brain, and blood-cancer derived cell lines, PcTF expression stimulates transcriptional activation of several anti-oncogenesis genes [48]. PcTF-mediated activation leads to the eventual loss of the silencing mark H3K27me3 and elevation of the active mark H3K4me3 at the tumor suppressor locus *CASZ1*.

To explore the therapeutic potential of fusion protein-mediated epigenetic interventions, we sought to investigate the behavior of PcTF in breast cancer cells lines that have been established as models for tumorigenesis [49–51]. Here, we extend our investigation of PcTF activity to three breast cancer-relevant cell lines. First, we investigated the transcription profiles of predicted PRC module genes in drug-responsive (MCF-7, BT-474) and unresponsive triple negative (BT-549) breast cancer cell lines. Receptor-negative BT-549 cells have a transcription profile and histology similar to aggressive tumor cells from patient samples [52,53]. Overexpression of PcTF in transfected breast cancer cells led to the upregulation of dozens of genes, including a common set of 19 genes in the interferon response pathway, as early as 24 hours after transfection. The transcriptome of BT-549 (triple-negative) showed the highest degree of PcTF-sensitivity. We observed that PcTF-sensitive genes are associated with a bivalent chromatin environment and moderate levels of basal transcription. Interestingly, these PcTF-sensitive genes do not overlap with very strongly repressed, PRC-enriched loci. This discovery provides new mechanistic insights into the state of genes that are poised for transcriptional activation via PcTF.

## RESULTS

### Differential regulation of genes in breast cancer cell lines

To determine expression levels of predicted PRC module genes, we profiled the transcriptomes of three breast cancer cell lines and the non-invasive, basal B cell line MCF10A [54,55] using next-generation deep sequencing of total RNA (RNA-seq). MCF7, BT-474, and BT-549 represent luminal A, luminal B, and basal B subtypes of breast cancer, respectively (Table 1) [49]. Previous studies have shown that gene expression profiles distinguish two major categories of cancer cell lines, luminal and basal, in patient-derived samples [56,57]. The basal class exhibits a stem-cell like expression profile [58], which is consistent with high levels of Polycomb-mediated repression at genes involved in development and differentiation [59,60]. Levels of the repressor protein EZH2 and the histone modification that it generates (H3K27me3) are elevated in MCF7, BT-474, and BT-549 compared to non-metastatic cells such as MCF10A (Table 1). A mechanistic link between Polycomb-mediated repression and tumor aggressiveness has been supported by a study where stimulation of the phosphoinositide 3-kinase (PI3K) signaling pathway, which induces a metastatic phenotype in MCF10A, is accompanied by increased H3K27me3 at several target genes [61,62]. We hypothesized that known Polycomb-repressed genes (the PRC module) would be down-regulated in the cancerous cell lines compared to MCF10A.

**Table 1.** Descriptions of the breast tissue-derived cell lines used in this study. ATCC = American Tissue Culture Center ID. Molecular subtype and marker expression status are from Neve et. al 2006 [49]: Estrogen receptor presence or absence (ER+/-), Progesterone receptor presence or absence (PR+/-), HER2 overexpression (HER2+), and TP53 mutation (*TP53*^*M*^). EZH2 and H3K27me3 were shown to be elevated compared to non-metastatic fibroblasts (a) [64], LNCaP (b) [63], MCF10A (c) [8,65,66], and HMEC (d).

| Cell line | ATCC | Sub-type | Markers [49] | EZH2 | H3K27me3 |
| --- | --- | --- | --- | --- | --- |
| MCF7 | HTB-22 | Luminal A | ER+, PR+ | Elevated<sup>a,b,c</sup> [63–65] | Elevated<sup>a</sup> [61,64] |
| BT-474 | HTB-20 | Luminal B | ER+, PR+, HER2+ | Elevated<sup>c</sup> [66] | Elevated<sup>d</sup> [61] |
| BT-549 | HTB-122 | Basal B, claudin-low | ER-, PR-, *TP53*<sup>M</sup> | Elevated<sup>c</sup> [8] | Elevated<sup>d</sup> [61] |
| MCF10A | CRL-10317 | Non-invasive/ Basal B | ER-, PR- | n/a | n/a |

Comparison of the expression profiles in untreated cells showed that the three breast cancer model cell lines were transcriptionally dissimilar to the control cell line MCF10A and that BT-549 and MCF7 were more similar to each other than either were to BT-474. Expression levels (FPKM values) across 63,286 gene protein coding transcripts (GRCh38 reference genome) were used to calculate Jensen-Shannon Divergence (JSD) (Methods and Fig. 2A). JSD values correspond to the similarity of the probability distributions of transcript levels for two RNA-seq experiments. Expression values for biological replicates showed the highest similarities (smallest distances) within cell types (Fig. 2A, upper grid). The largest distances were observed between MCF10A and the three cancer cell types: 0.461 for BT-549, 0.476 for MCF7, and 0.511 for BT-474 (Fig. 2A, lower grid). A similarly high JS distance was observed for BT-549 versus BT-474 (JSD = 0.464), suggesting that these cancer cell lines are transcriptionally distinct. BT-549 and MCF7 showed the highest similarity, with a cumulative JSD of 0.357. This observation contrasts with other reports where BT-549 and MCF7 are described as transcriptionally and phenotypically different [54,67]. Differences in transcription profiling methods, RNA-seq used here and the DNA oligomer microarray chip used by others, may underlie the different outcomes.

Differential expression between cell lines for individual genes (Fig. S1) followed similar trends as those observed for the global JSD analysis. We used an expression comparison algorithm (Cuffdiff [69]) to identify genes that were differentially expressed (2-fold or greater difference in expression, *q* value ≤ 0.05) or similarly expressed (less than 2-fold difference, *q* value ≤ 0.05) between cell types. Comparisons that included MCF10A showed the highest numbers of differentially-expressed genes, as well as the lowest numbers of similarly expressed genes. This result further supports transcriptional differences between the cancerous cell lines and MCF10A (Fig. S1).

Next, we determined expression levels within groups of predicted PRC-regulated genes and observed that expression within these subsets is lower in the three cancer cell types than in MCF10A. We used data from other breast cancer cell line studies of MCF7 and MDA-MB-231 to classify a subset of PRC target genes based on H3K27me3 enrichment or binding of EZH2, an enzyme that generates the H3K27me3 mark (see Methods). Only 245 gene IDs were shared between the H3K27me3 and EZH2 subsets. Although these two groups are mostly distinct, both showed low median expression values (FPKM < 2), which suggests epigenetic repression (Fig. 2B). Median expression levels of predicted PRC module genes were reduced in the cancer cell lines compared to the non-cancer cell line. The H3K27me3-marked subset showed median log10(FPKM) values for BT-474 (−1.66), MCF7 (−1.16), and BT-549 (−1.15) that were slightly lower than MCF10A (−1.10) (Fig. 2B, middle plot). The median FPKM values for EZH2 targets were dramatically lower (zero signal) in the cancer cell lines, while the median value was higher (−1.65) for MCF10A (Fig. 2B, right). Overall, H3K27me3 and EZH2 enrichments from two breast cancer cell lines (MCF7 and MDA-MB-231) correspond to relatively low expression in all three breast cancer cell lines studied here. This result is consistent with the roles of H3K27me3 and EZH2 in cancer-associated gene silencing.

**Figure 2.**
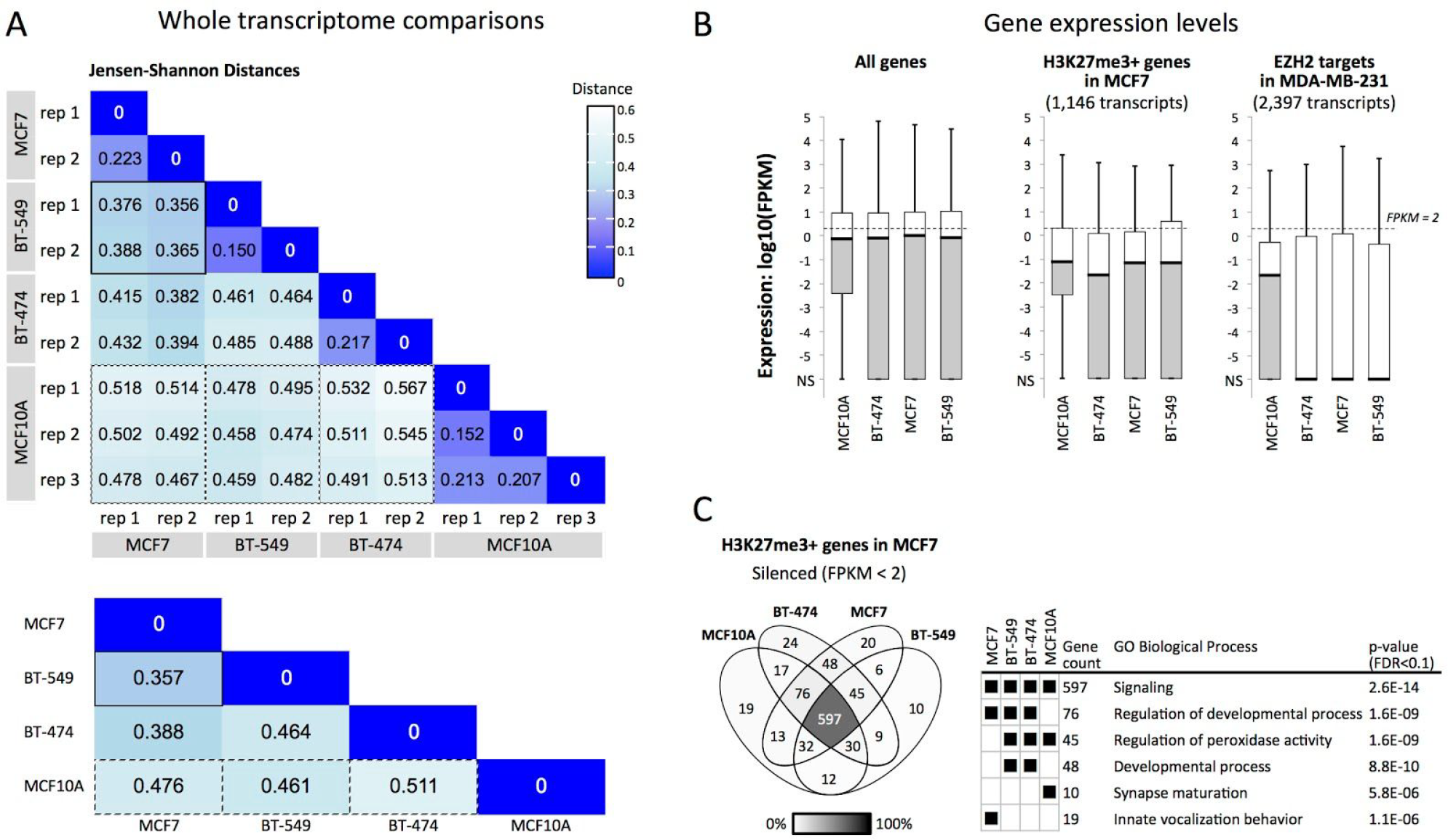
Comparisons of transcription profiles of three model breast cancer lines (MCF7, BT-549, BT-474) and a control non-cancer line (MCF10A). (A) Jensen-Shannon Divergence (JSD) values were calculated as the similarity of the probability distributions of expression levels (FPKM values) for 63,286 total transcripts, which include 22,267 protein-coding transcripts. In the lower grid, cummeRbund [68] was used to consolidate replicates and to calculate overall JSD between cell types. Solid border, BT-549 vs. MCF7, smallest JSD; dashed border, JSD’s for MCF10A vs. cancer cell lines. (B) The boxplots show gene expression values (center line, median; lower and upper boxes, 25th and 75th percentiles; lower and upper whiskers, minimum and maximum) for all protein-coding transcripts (22,267), H3K27me3-positive (1,146) or EZH2-positive (2,397) protein-coding loci. NS, no signal. (C) The Venn diagram includes HGNC symbols of genes that are H3K27me3-positive (middle box plot, panel B) and are silenced (FPKM < 2) in at least one cell type. GO term enrichment *p*-values are shown only for subsets where FDR < 0.1.

To determine whether individual predicted PRC target genes were similarly regulated across cell lines, we compared two groups of genes that were categorized by expression level: silenced (FPKM < 2) [70,71] or expressed (FPKM ≥ 2) (Fig. S2). In each cell type, genes with silenced expression levels included 70.2% - 79.3% of the H3K27me3-marked loci (Fig. S2) and 78.4% - 82.2% of the EZH2-enriched loci. About one quarter of the genes (17.8% - 29.8%) showed some expression (FPKM ≥ 2) and only 16.7% - 8.2% were expressed at FPKM ≥ 10. The set of 45 H3K27me3-enriched repressed genes shared by the three cancer cell lines BT-474, BT-549, and MCF7 (Table S1) shows strong representation of the gene ontology processes “regulation of peroxidase activity” (GOrilla[72], *p* = 5.84E-6, FDR = 8.85E-2; Fig. 2C) and “ectoderm development” (Panther[73], *p* = 1.07E-4, FDR = 2.61E-2). The silencing of lipoxygenase (*ALOXE3*) and and inhibitor of peroxidase (*LRRK2*) may contribute to elevated pro-cancer COX-mediated peroxidase activity [74,75]. Low levels of *ALOXE3*, *ADRB2*, *BNC1*, *BTC*, *CCNO*, *ETV4, MCIDAS*, *PID1*, *SPRR2D*, and *ZBTB16* are consistent with the epigenetic repression of pro-differentiation pathways in cancer cells. We hypothesized that these PRC-module genes would become activated in the presence of the synthetic regulator PcTF, which interacts with the repressive H3K27me3 mark.

### PcTF-sensitive interferon response genes are shared across three cancer cell types

We investigated changes in the transcriptomes of PcTF-expressing breast cancer cells over time. We transfected cells with PcTF-encoding plasmid DNA (previously described [48]) and allowed them to grow for 24, 48, and 72 hours before extracting total RNA for sequencing. RNA-seq reads were aligned to a human reference genome GRCh38 that included the coding region for PcTF (see Methods). No reads aligned to the PcTF coding sequence in control, untransfected cells. In the transfected cells, PcTF expression levels were highest at 24 hours and decreased 1.6 to 5.5-fold every 24 hours (Fig. 3A). We observed a similar trend with other cancer cell lines in a previous study [48]. One outlier sample, a replicate for BT-474 cells expressing PcTF for 48 hours, had a markedly different PcTF expression level (Fig. 3A) and genome-wide transcription profile (Fig. S3) and was therefore omitted from further analyses.

**Figure 3.**
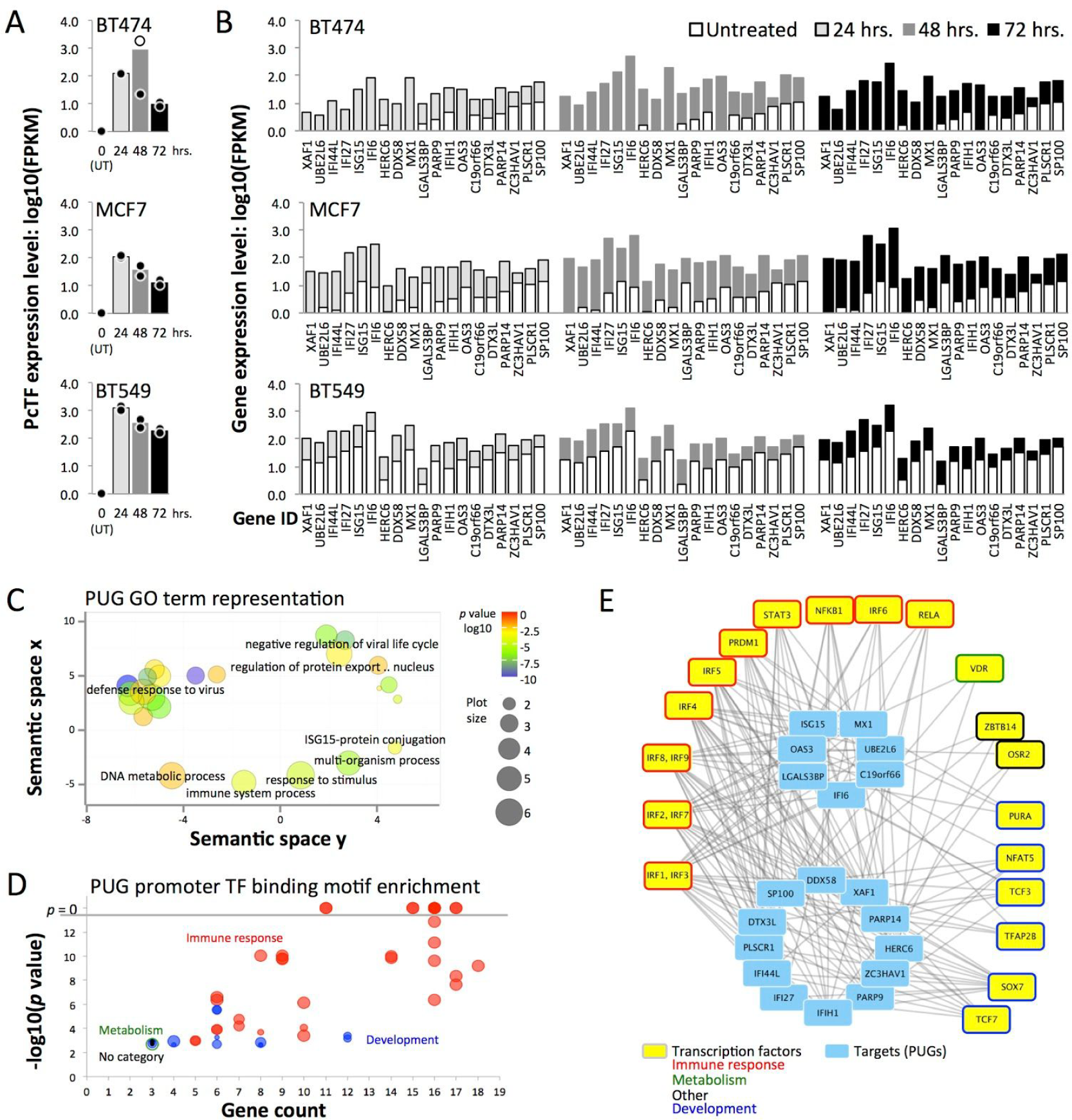
PcTF-expressing breast tissue-derived cell lines show upregulation of interferon (IFN) pathway genes. (A) Charts show log10(FPKM) of PcTF for untransfected cells (UT) and at 24, 48, and 72 hours following transfection of each cell line. The outlier for BT-474 (48 hrs, replicate 1) was omitted from subsequent analyses. Dots, each replicate library; bars, mean of values from the two replicates. (B) Mean log10(FPKM) values are shown for 19 Polycomb-upregulated genes genes (PUGs; FC ≥ 2, *q ≤* 0.05 at all time points in all three cell lines), sorted from lowest to highest average expression level in untreated cells. (C) Gene ontology (GO) Biological Process term enrichment for the 19 PUGs is represented the bubble chart. GO clusters and representative terms (black labels) are plotted based on semantic similarities in the underlying GOA database. (D) Overrepresentation of transcription factor (TF) binding motifs [76] at the promoters of PUGs (*p*-value < 0.05/19.0, Bonferroni correction). (E) Transcription factors (outermost boxes) associated with promoter motifs from panel D are shown in the network graph.

Nineteen genes were upregulated at least 2-fold (*q* value ≤ 0.05) at all time points in all three cell lines (Fig. 3B): *C19orf66*, *DDX58, DTX3L, HERC6, IFI27, IFI44L, IFI6, IFIH1, ISG15, LGALS3BP, MX1, OAS1, OAS3, PARP9, PARP14, PLSCR1, SP100, UBE2L6*, and *XAF1*. Here, we refer to this subset PcTF-upregulated genes, or PUGs. The most significantly enriched GO terms for this set include “defense response to virus” and “negative regulation of viral life cycle” (Fig. 3C). An investigation of regulator motif enrichment at the promoters of PUGs revealed significant overrepresentation of transcription factors involved in immune response and tissue development processes (Fig. 3D). Fifteen of the 22 transcription factors showed detectable levels of expression in all three cell lines (Fig. S4). *IRF1*, *IRF7*, *IRF9*, and *PRDM1* showed significant upregulation (FC ≥ 2, q ≤ 0.05) in PcTF-expressing cells. Promoter motifs for *IRF1* and *IRF3* were present at all 19 PUGs (Fig. 3E). Therefore, regulation of PUGs may be driven by PcTF-mediated activation of IRF1.

Different subsets of genes were up- or down-regulated at least two-fold (*q* value ≤ 0.05) early, late, or across all time points during PcTF expression (Fig. 4). Of the genes that showed at least a two-fold change in either direction, the vast majority were up-regulated (Fig. 4A). We also observed that depending on the cell line, two or three predicted regulators of PUGs, including *IRF1*, *IRF7*, *IRF9*, and *PRDM1*, became significantly upregulated (Fig. 4B). This result suggests that the IFN response might be mediated through upregulation of master regulators. Thus, PcTF may target silenced chromatin at *IRF1*, *IRF7*, *IRF9*, and *PRDM1* and not necessarily at PUGs.

**Figure 4.**
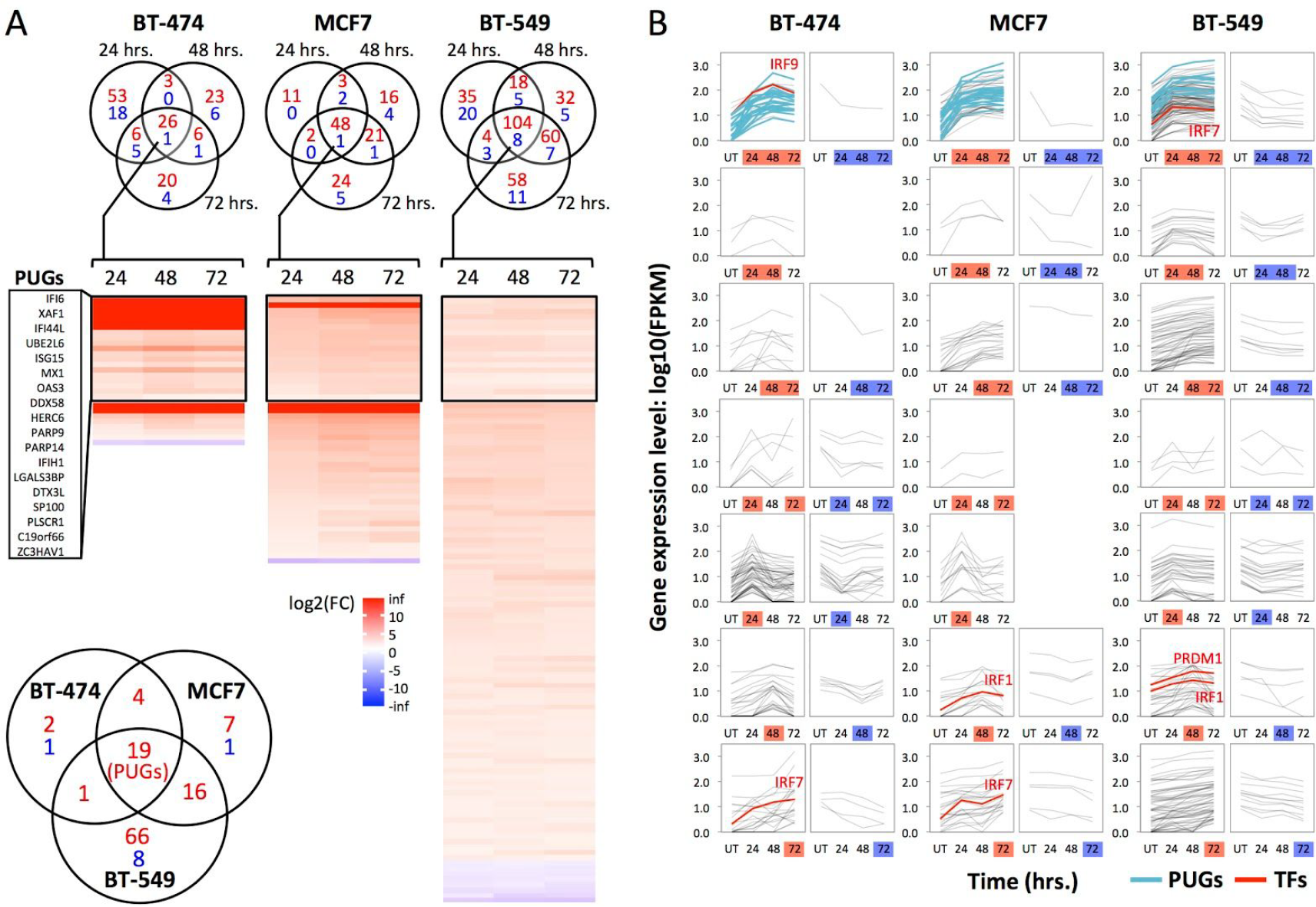
PcTF-sensitive genes include cell-type specific groups in addition to PUGs. (A) The Venn diagrams show genes with expression levels that changed at least 2-fold in either direction (*q* value ≤ 0.05) at one or more time points in PcTF-expressing cells versus untransfected cells. Red, up-regulated; blue, down-regulated. The heat maps show fold-change (log2(FC)) values for genes that significantly changed (*q ≤* 0.05) at all three time points (center regions of the Venn diagrams). The lower left Venn diagram compares these genes between cell types. (B) Expression profiles (log10(FPKM)) of cells before (UT, untransfected), and 24, 48, or 72 hours after PcTF transfection for all genes with expression levels that changed at least 2-fold in either direction (see Venn diagrams in panel A).

Our results also show that the PcTF-activated genes had virtually no overlap with the 45 H3K27me3-enriched, silenced genes (FPKM < 2) shared by the three cancer cell lines (Fig. 2C, Table S1). Only one of these 45 genes, *PID1*, became upregulated in any cell line (BT-549 at 48 and 72 hours). In this study we observed that the genes that were up-regulated came from the pool of low- to moderate-expressing genes. So far, our results suggest that PcTF-mediated activation requires a moderate level of basal expression at the target gene. This idea may be counterintuitive since H3K27me3 mark, the target of PcTF [47], is essential for transcriptional repression according to the model for Polycomb-mediated regulation, which is supported by a wealth of data [77]. However, a recent study using genome-wide ChIP-seq and transcription profiles in murine cells showed that H3K27me3 was enriched at genes with low levels of expression and depleted at completely silenced genes, and highly expressed genes[78]. We were prompted to investigate whether the chromatin features at PcTF-activated genes might reflect a low to moderate expression state.

### PcTF-sensitive loci bear repression- and activation-associated chromatin features

To investigate the contribution of local chromatin states to PcTF-mediated gene regulation, we analyzed histone modifications and RNA polymerase II enrichment at PcTF-upregulated genes in MCF7. Here, we utilized the extensive public ChIP-seq data that is available for the MCF7 cell line to investigate chromatin features. The 125 genes that were significantly upregulated (FC ≥ 2, *q ≤* 0.05) at one, two, or all time points in MCF7 (see Fig. 4B) showed a range of H3K27me3 mean enrichment values across 10 kb centered around each transcriptional start site (Fig. 5A). Consistent with PUGs, the 106 additional upregulated genes showed significant overrepresentation of interferon response-related processes (GO biological process “type I interferon signaling pathway,” *p* = 4.08E-28, FDR = 6.21E-24).

**Figure 5.**
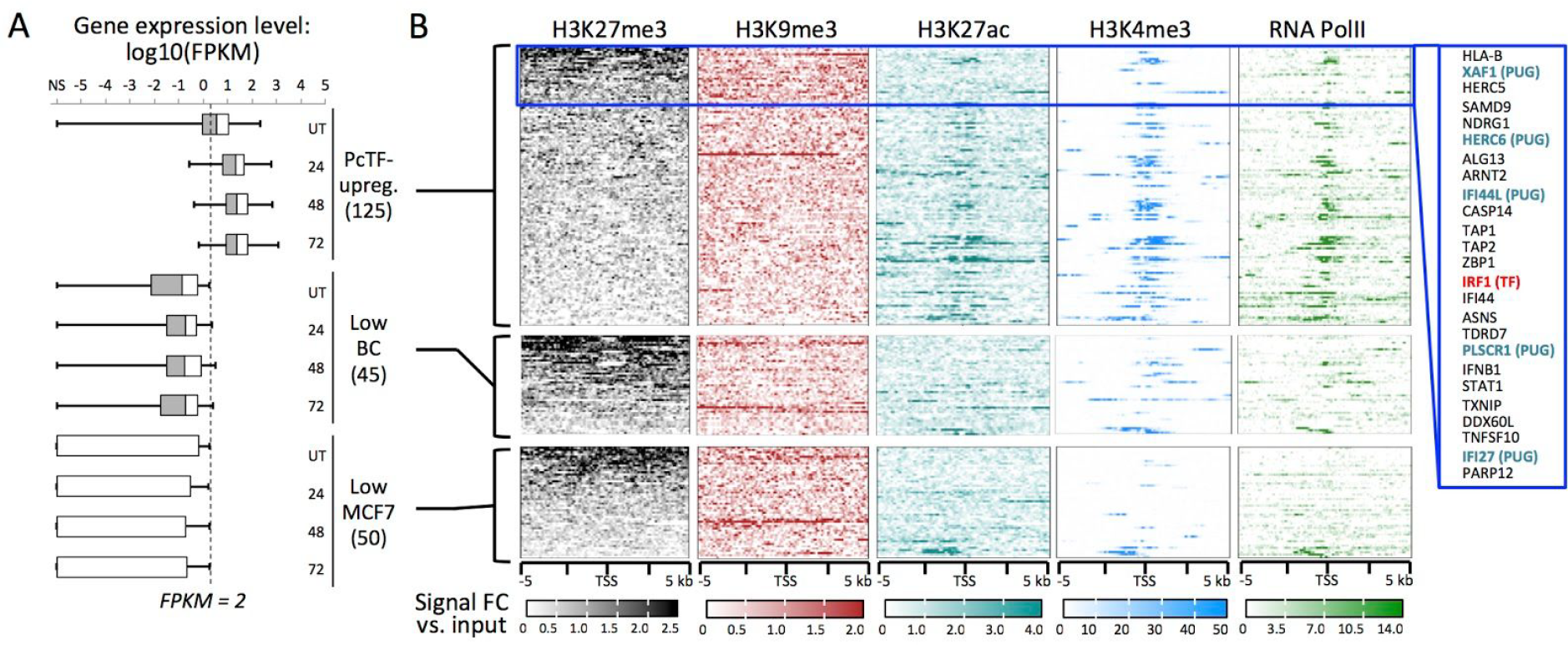
Comparison of chromatin features at PcTF-activated and non-activated genes in MCF7. (A) Box plots show expression levels (center line, median; left and right boxes, 25th and 75th percentiles; left and right whiskers, minimum and maximum) in untreated and PcTF-treated cells (24, 48, and 72 hrs) for each of the following gene subsets: PcTF-upreg., 125 genes that are upregulated (FC ≥ 2, *p ≤* 0.05) in MCF7-expressing cells at one or more time points; Low BC, 45 H3K27me3-enriched genes that are repressed (FPKM < 2) in all three cancer cell lines (see Fig. 2C); Low MCF7, 50 genes that are repressed (FPKM < 2) in MCF7. TSS plots show ChIP signals of silencing-associated (H3K27me3, H3K9me3) and activation-associated (H3K27ac, H3K4me3) histone modifications, as well as RNA Polymerase II. Genes within the top 20% of mean values for H3K27me3-enrichment (within 10 kb) are highlighted (blue box).

Genes within the highest 20% of mean values for H3K27me3 included the predicted regulator *IRF1* (Fig. 2D, E) and 5 of the 19 PUGs. Other PcTF-responsive genes that lack the H3K27 methylation mark might represent downstream targets of the products expressed from targets of PcTF. Mean enrichments of H3K9me3 (Fig. 5A), a modification that is frequently found at constitutive pericentric heterochromatin and non-coding DNA [79–81], showed no pattern that resembled H3K27me3. PcTF-responsive genes tended to be distributed along chromosome arms rather than concentrated near centromeres (Fig. S4). This suggests that PcTF target sites coincide more closely with the distribution of facultative chromatin and epigenetically-regulated cell development genes [59,82].

Enrichments for the features associated with active expression, H3K27ac, H3K4me3, and RNA Pol II were stronger at PcTF-responsive genes than at PcTF non-responsive genes (Fig. 5B). Regions containing PcTF-activated genes include interspersed peaks of H3K27me3 and H3K4me3 (Fig. S5), which is characteristic of bivalent domains that are poised for activation [15,83]. We conclude that under the conditions tested here, strongly repressed genes are resistant to PcTF-mediated activation while an intermediate regulatory state, where silent and active marks are present, supports PcTF activity.

Two substantially different mechanisms might account for the results observed so far. First, target gene activation may depend upon PcTF’s interaction with and disruption of silenced chromatin. In previous work, we established that PcTF activity requires the histone-binding PCD domain [48,84] and the presence of H3K27me3 near the target gene [84] to disrupt epigenetic silencing. Work reported by others demonstrated activation of interferon networks through the disruption of chromatin-mediated repression with small molecule inhibitors. Treatment of breast cancer cell lines (including BT-474 and MCF7) with DNA methyltransferase (5-azacitidine) led to activation of *DDX58*, *IFI27*, *IFI6*, *IFIH1*, *ISG15*, *MX1*, *OAS3*, *UBE2L6*, *XAF1* (9 of the 19 PUGs), and other genes [33]. Furthermore, inhibitors of histone deacetylase, a class of enzymes that support repressed chromatin, stimulate rapid activation of interferon (IFN) genes in human and mouse cells [85].

Second, introduction of foreign nucleic acids into the cells could have indirectly stimulated the interferon response via sequence non-specific effects [86–90] without interaction of PcTF with chromatin. Microarray-based transcriptome profiling of MCF7 cells transfected with Lipofectamine-pM1-MT vector complexes showed upregulation of *HERC6*, *IFIH1*, *ISG15*, *LGALS3BP, MX1*, *OAS3, PLSCR1,* and *UBE2L6* [89], which represent 8 of the 19 PUGs. Small RNA-induced knockdown of GAPDH in renal carcinoma cells was accompanied by increased expression of *IFI6*, *OAS3*, and *UBE2L6* [86]. *MX1, IRF1 and IRF7* became activated following electroporation (nucleofection) of NIH3T3 cells with control empty plasmids pcDNA3.1 (the origin of the plasmids used in our study), phGF, and pEGFP-N1 [90]. To investigate nonspecific effects from foreign nucleic acids, we used reverse transcription followed by quantitative PCR to measure expression levels of PcTF-responsive genes in cells that expressed a truncated version of PcTF as a control, as described in the following section.

### Foreign RNA from a PcTF-deletion mutant is insufficient for sustained expression of *XAF1* in MCF7

We asked whether the presence of the PcTF transgene and its transcribed RNA were responsible for the consistent interferon response in breast cancer cells. Using transient transfections, we had established that PcTF-mediated activation of genes could be detected over background at multiple time points. However, in this experiment PcTF levels decreased over time (Fig. 3A), which prevents us from distinguishing time-versus dose-dependent effects on gene regulation.

Therefore, we constructed stable transgenic cell lines to enable constant expression of the fusion protein over time. We were able to generate viable, transgenic lines from MCF7 cells. Expression of PcTF or a control fusion protein that lacks the histone-binding domain (Pc_Δ_TF) was placed under the control of the rtTA activator, which binds to the *pTet* promoter in the presence of doxycycline (dox) (Fig. 6A). Expression of rtTA was indicated by constitutive GFP expression, and inducible nuclear localization sequence-tagged PcTF was detected as an RFP signal after treatment with doxycycline (Fig. 6B). We used reverse transcription followed by quantitative PCR (RT-qPCR) to measure the expression levels of PcTF and a subset of PcTF-sensitive genes that were identified in the RNA-seq experiment.

**Figure 6.**
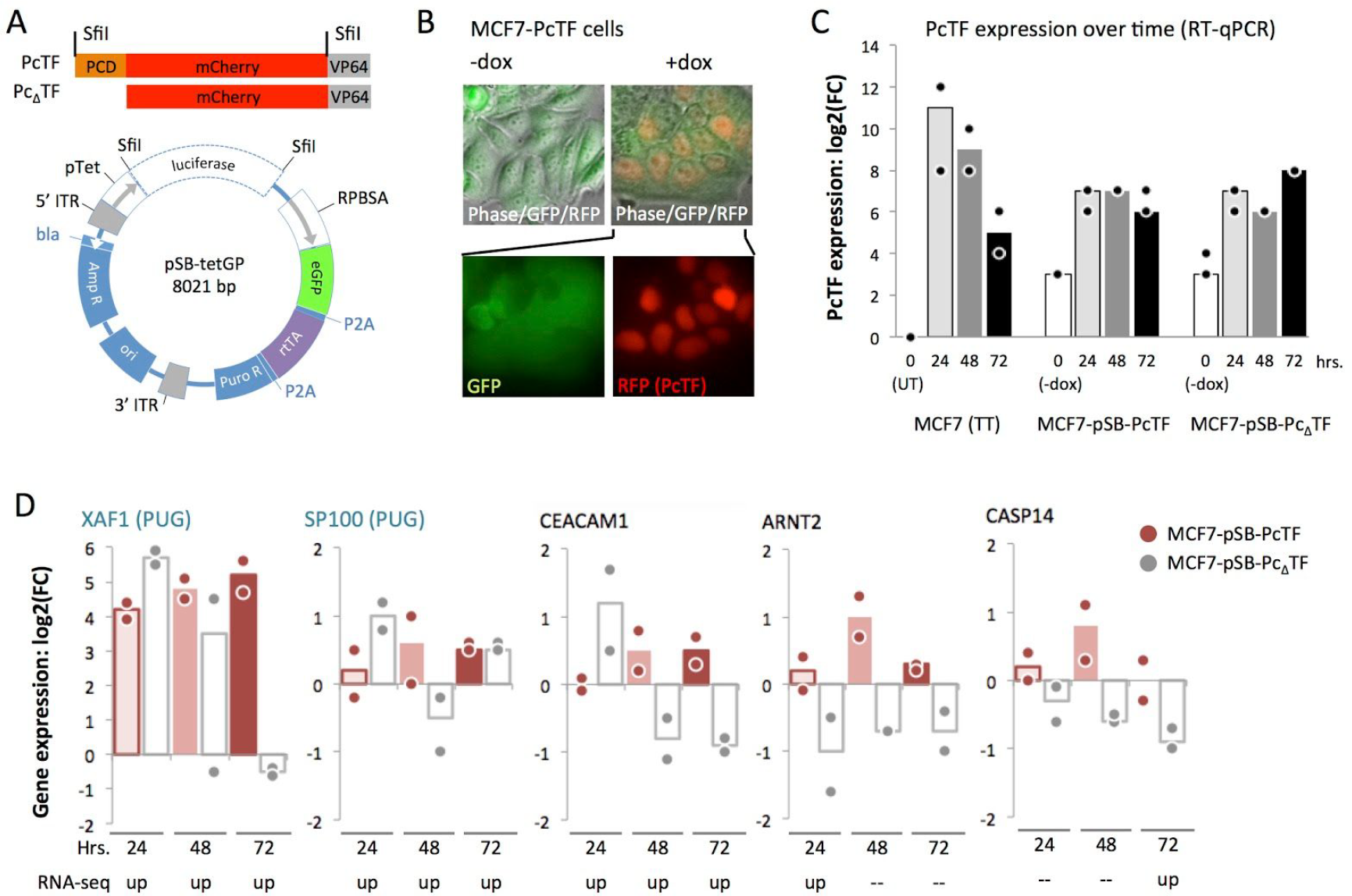
RT-qPCR analysis of gene expression in stable, transgenic PcTF-expressing cells. (A) *SfiI*-flanked PcTF or PcΔTF constructs (top) were cloned into the pSBtet-GP expression vector (bottom), resulting in the replacement of the *luciferase* reporter with fusion protein ORFs. (B) Fluorescence microscopy of the MCF7-PcTF transgenic cell line. (C) Time course qRT-PCR for PcTF. (D) Time course qRT-PCR for select genes. For all RT-qPCR experiments *n* = two cDNA libraries from independent transfections or dox treatments. FC, fold change relative to “no dox” controls, calculated as double delta C_p_ (see Methods).

RT-qPCR using a universal mCherry-specific primer set confirmed that PcTF expression levels decreased over time in transiently transfected cells (Fig. 6C) as observed for FPKM values from the RNA-seq experiment (Fig. 3A). The stable transgenic cells showed low levels of fusion protein mRNA in the initial uninduced (-dox) state compared to untransfected MCF7 cells. Exposure to 1 μg/mL dox increased PcTF and Pc_Δ_TF levels by an order of magnitude. These levels were slightly higher than the PcTF expression levels observed in transiently transfected cells at the 72-hour time point, and remained relatively constant over time. Fold-change (compared to untransfected cells) remained within values of 67 - 192 at 24, 48, and 72 hours.

For RT-qPCR analysis of PcTF-sensitive targets, we were able to design and validate specific assays for a subset of genes that were significantly upregulated at one or more time points in MCF7, including two PUGs (*XAF1*, *SP100*) and others. *XAF1* was the most strongly upregulated across all three time points (18 to 36-fold) (Fig. 6D). The other five genes showed slight upregulation in response to dox-induced PcTF expression. The weaker response of these genes compared to *XAF1* could be explained by a smaller dynamic range, where there is little difference between the basal versus activated expression level. Furthermore, these genes may have been slightly upregulated prior to dox treatment since PcTF was detected at low levels before induction (Fig. 5C).

At the 24 hour time point, *XAF1*, *SP100*, and *CEACAM1* became up-regulated in truncation-expressing cells, suggesting an initial nonspecific response to transgenic PcΔTF RNA. At 48 and 72 hours, gene expression decreased in the presence of Pc_Δ_TF. Over time, expression remained upregulated in the presence of PcTF compared to Pc_Δ_TF at *XAF1*, *CEACAM1*, and *ARNT2*. Overall, these results suggest that for certain genes (*XAF1*, *CEACAM1*, and *ARNT2*), maintenance of the PcTF-induced activated state requires interaction with chromatin through the H3K27me3-binding PCD motif.

### Tumor suppressor and BRCA pathway genes become upregulated in PcTF-expressing cells

To explore the clinical implications of PcTF-mediated transcriptional regulation, we determined the representation of known tumor suppressor genes amongst PcTF-responsive loci. For this analysis we used a tumor suppressor gene set that includes 983 candidate anti-cancer targets that are down-regulated in tumor samples (Methods). Of these, 589 include BRCA human tumor suppressor genes (TSGs) that are repressed in invasive carcinoma samples compared to normal tissue samples [91,92]. The genes were classified as tumor suppressors based on text-mining of cancer research literature, and manual assessment of relevant cancer types and molecular pathways (TSGene 2.0) [91,92].

To identify TSGs that are upregulated in response to PcTF, we compared the upregulated subset (FC ≥ 2, *q ≤* 0.05) to the 983 candidate anti-cancer genes identified by TSGene 2.0. Fifteen of the 983 TSGs were upregulated across all three time points in at least one of the cell lines (Fig. 7A). Information from genecards.org [93] further validated the association of these 15 genes with tumor suppressor activity. Of the fifteen upregulated TSGs, seven belong to the breast cancer susceptibility (BRCA) pathway: *CDKN1A*, *PML*, *ANGPTL4*, *CEACAM1*, *BMP2*, *SP100*, *TFPI2*.

**Figure 7.**
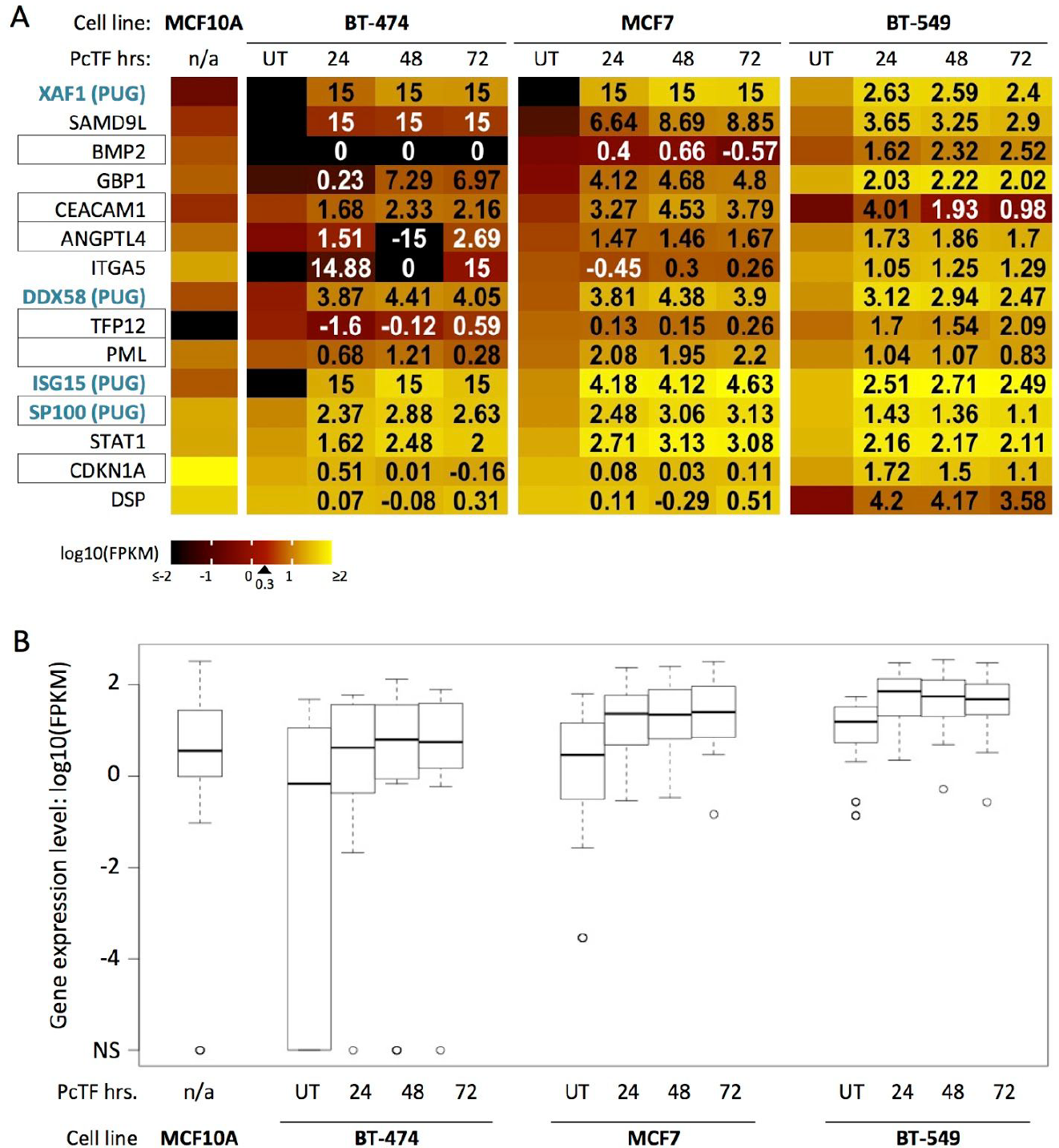
Tumor suppressor genes show increased expression in PcTF-expressing cancer cell lines. (A) Individual log10(FPKM) (color scale) for each of the tumor suppressor genes in A. Black boxes highlight BRCA pathway genes. Genes are sorted from lowest to highest expression in untreated MCF7 cells. Numbers in the PcTF-treatment columns show log2 fold change values compared to UT. 15, infinite positive fold change where no expression was detected in untreated cells; −15, infinite negative fold-change where no expression was detected in treated cells. (B) Box plots show expression values (center line, median; lower and upper boxes, 25th and 75th percentiles; lower and upper whiskers, minimum and maximum) across three time points (24, 48, and 72 hours) for fifteen tumor suppressor genes where upregulation was at last two-fold (*q ≤* 0.05) relative to the untreated control (UT) in at least one of the cell lines.

Cell line comparisons of RNA-seq FPKM values for the fifteen tumor suppressor genes showed that median expression was lower in untreated BT-474 and MCF7 than in the non-cancerous MCF10A cell line (Fig. 7B). This result is consistent with the idea that epigenetic repression of TSGs supports a cancerous cell phenotype. In PcTF-expressing cells, the median expression of the fifteen tumor suppressor genes was increased at all time points compared to the untreated samples for each cancer cell line (Fig. 7B). Interestingly, the median FPKM value for the 15 TSGs was higher in BT-549 than in MCF10A. Closer examination of the the individual genes revealed that expression levels for *BMP2*, *CEACAM1*, *CDKN1A*, *DSP* are lower in BT-549 than in MCF10A (Fig. 7A). These genes become upregulated in PcTF-expressing cells. These results demonstrate that PcTF stimulates conversion of the expression state of several tumor suppressor genes from silenced to active.

## DISCUSSION

As the importance of global chromatin-mediated dysregulation in oncogenesis is coming to light, scientists are becoming more interested in using inhibitors to block master regulators of repressive chromatin (i.e., HDACs, DNMTs, HMTs[18,28,33,34,36]) to investigate and treat cancer. This approach has been recently described as “macrogenomic engineering”[94]. A key advantage of broad epigenetic manipulation is that it is DNA sequence-agnostic; the therapeutic effect potentially does not require *a priori* knowledge of patient-specific sequence variations at a candidate target gene or genes. Cancer tissues often accumulate extensive DNA lesions, from small insertions and deletions to large chromosome rearrangements. Therefore, editing or activating single targets may not be effective in some cells. In this report we present a synthetic approach to macrogenomic engineering, a fusion protein that physically bridges a chromatin feature at silenced genes (H3K27me3) with proteins that drive gene activation. Our previous studies have established that PcTF specifically interacts with H3K27me3 *in vitro* [47], and drives the activation of hundreds of repressed loci including master regulators and tumor suppressors in bone, blood, and brain cancer derived model cell lines [48]. In our current report, we discovered a core set of interferon-pathway-related genes that responded to PcTF in three distinct breast cancer cell lines.

Several factors can contribute to transcriptomic variations in breast cancer subtypes, such as differences in the abundance of wild type or mutated transcription factors, mutations that impact the stability and turnover of RNA transcripts, and dysregulation of histone-modifying enzymes [13]. It is important to determine the relationship between phenotypic subclasses and transcription profiles [16,54,67,95] to elucidate cancer mechanisms and drug targets for more effective treatments. Establishing a link between transcriptomes and phenotypes may require further research. We observed that the transcription profile of BT-549 (invasive basal B) is more similar to MCF7 (luminal) than either were to BT-474 (luminal). In contrast, other reports have shown clear distinctions between the transcription profiles and phenotypes of BT-549 and MCF7 [54,67]. Differences in transcript profiling methods, our RNA-seq and JSD analysis versus the DNA oligomer arrays used by others, may account for this conflicting result. Further, we acknowledge that the JSD may be driven by a few genes with high expression and high variance, which could account for some of the patterns.

Diversity of breast cancer cell transcriptomes poses a formidable challenge for the development of drugs that target specific proteins, genes, and pathways. Our results demonstrate that activation of a common set of genes can be achieved by direct targeting of H3K27me3 with a fusion activator (PcTF) in three distinct model breast cancer cell lines that show distinct basal gene-expression levels. The 19 common PcTF-upregulated genes (PUGs) show significant overrepresentation of the GO biological processes “defense response to virus” and “negative regulation of viral life cycle.” A larger set of 125 genes that are upregulated at any time point in MCF7 (Fig. 4, 5) are associated with “type I interferon signaling pathway”. Enrichments of H3K27me3 signals near the promoters of five PUGs (*XAF1*, *HERC6*, *IFI44L*, *PLSCR1*, *IFI27*) and a predicted regulator of all 19 PUGs (*IRF1*), suggest that PcTF accumulates near these promoters and recruits transcriptional activation machinery as demonstrated for *CASZ1* in a previous study[48]. Another potential mechanism for stimulation of the IFN pathway is epigenetic de-repression of endogenous retroviral dsRNA production, as observed during treatments with inhibitors against DNA methyltransferases histone deacetylases [96–98]. It has been proposed that this process mimics a viral infection that makes the cancer cell a target for destruction by the immune system or immunotherapies [99].

While many H3K27me3-enriched genes were upregulated in MCF7, many were non-responsive under the conditions tested here (up to 72 hours of PcTF expression). At PcTF-responsive genes, levels of H3K4me3 and H3K27ac were higher than at silenced non-responsive genes. Therefore, the chromatin at PcTF-responsive genes may support a low or intermediate expression state. Berrozpe et al. recently reported that Polycomb complexes preferentially accumulate at weakly expressed genes rather than strongly silenced or strongly expressed genes [78]. In our experiments, specific PRC-regulated genes may have been expressed at low to intermediate levels and then further upregulated upon exposure to PcTF. Our analysis of PcTF-regulated genes and chromatin states paves the way for future studies to further resolve chromatin features that distinguish regulatable PRC-repressed genes in cancer cells.

So far, low molecular weight compounds are the predominant method for epigenetic research and interventions. Their ease of delivery, orally or intravenously, make these compounds a very attractive approach for *in vivo* studies and cancer treatment. However, small compounds have a very limited range of biological activity, *e.g.* as ligands for specific proteins, compared to macromolecules. Transgenic and synthetic transcription factors expand the repertoire of epigenetic drug activity by allowing selective control of therapeutic genes in cancer cells [102–105]. Protein expression often relies on inefficient and possibly mutagenic nucleic acid delivery, which poses a significant barrier for many potential synthetic biologics. Recent advances in large molecule carriers such as cell penetrating peptides [106–108] provide a positive outlook for cellular delivery of purified proteins.

## CONCLUSIONS

In conclusion, we have demonstrated that PcTF stimulates broad changes in expression, reminiscent of the effects observed for small-molecule epigenetic drugs, that could disrupt the immune evasion phenotype of cancer. Activation of IFN pathways genes has important implications for cancer research and therapy. Other studies have linked high levels of expression from interferon pathway genes with a non-cancerous phenotype. In breast cancer, expression of an immune response gene subgroup, which includes *ISG15*, *MX1*, and other interferon genes, has been associated with improved prognosis in triple negative breast cancers [109,110]. It will be eventually important to determine if PcTF proteins meet or exceed the efficacy of low molecular weight epigenetic drugs in tumor and patient-derived models. At present, PcTF an its variants [47] represent a new exploration space for rationally-designed epigenetic interventions.

## MATERIALS AND METHODS

### DNA constructs

Plasmids were constructed to express fusion proteins either constitutively or in the presence of doxycycline. The plasmid for constitutive expression of PcTF, hPCD-TF_MV2 (KAH126), was constructed as previously described [84]. The doxycycline-inducible transgene PcTF_pSBtet-GP was constructed by ligating 50 ng of PCR amplified, SfiI-digested PcTF fragment with a SfiI-linearized pSBtet-GP vector [111] (Addgene #60495) at a ratio of 5 insert to 1 vector in a 10 uL reaction (1 uL 10x buffer, 1 uL T4 ligase). The same procedure was used to build constructs for dox-inducible Pc_Δ_TF expression. Primers used for the PCR amplification step are as follows: Forward 5’-tgaaGGCCTCTGAGGCC<u>aattcgcggccgcatctaga</u>, Reverse 5’-gcttGGCCTGACAGGC<u>tgcagcggccgctactagt</u>. Template-binding sequences are underscored. Adjacent nucleotides were designed to add *SfiI* restriction sites (uppercase) to each end. The full annotated sequences of all plasmids reported here are available online at Benchling - Hayneslab: Synthetic Chromatin Actuators (https://benchling.com/hayneslab/f/S0I0WLoRFK-synthetic-chromatin-actuators/).

### Cell culture and transfection

MCF7 (ATCC HTB-22) cells were cultured in Eagle’s Minimal Essential Medium supplemented with 0.01 mg/mL human recombinant insulin, 10% fetal bovine serum, and 1% penicillin and streptomyicn. BT-474 cells (ATCC HTB-20) were cultured in ATCC Hybri-Care Medium supplemented with 1.5 g/L sodium bicarbonate, 10% fetal bovine serum, and 1% penicillin and streptomycin. BT-549 cells (ATCC HTB-122) were cultured in RPMI-1640 Medium supplemented with 0.0008 mg/mL human recombinant insulin, 10% fetal bovine serum, and 1% penicillin and streptomycin. MCF-10A cells (ATCC CRL-10317) were cultured in Mammary Epithelial Cell Growth Medium (Mammary Epithelial Cell Basal Medium and BulletKit supplements, except gentamycin-amphotericin B mix), supplemented with 100 ng/mL cholera toxin. Cells were grown at 37 °C in a humidified CO_2_ incubator. PcTF-expressing MCF7, BT-474, and BT-549 cells were generated by transfecting 5×10^5^ cells in 6-well plates with DNA/Lipofectamine complexes: 2 μg of hPCD-TF_MV2 plasmid DNA, 7.5 μl of Lipofectamine LTX (Invitrogen), 2.5 PLUS reagent, 570 µl OptiMEM. Control cells were mock-transfected with DNA-free water. Transfected cells were grown in pen/strep-free growth medium for 18 hrs. The transfection medium was replaced with fresh, pen/strep-supplemented medium and cells were grown for up to 72 hrs.

### Generation of stable cell lines

To generate doxycycline-inducible cell lines, MCF7 cells were transfected with the transposase-expressing plasmid SB100X and either hPCD-TF_pSBtet-GP or TF_pSBtet-GP (19:1 molar ratio of pSB to SB100X), under the same conditions as described above. After 24 hrs, the transfection medium was replaced with fresh, puromycin-supplemented medium (0.5 μg/mL). Cells were then grown until cell cultures were >90% GFP-positive as measured by flow cytometry. Total culture time was 2-3 weeks per cell line.

### Preparation of total mRNA

Total messenger RNA was extracted from ~90% confluent cells (~1-2×10^6^). Adherent cells were lysed directly in culture plates with 500 μl TRIzol. TRIzol cell lysates were extracted with 100 μl chloroform and centrifuged at 12,000 xg for 15 min. at 4°C. RNA was column-purified from the aqueous phase (Qiagen RNeasy Mini kit 74104).

### Quantitative reverse transcription PCR (qRT-PCR)

SuperScript III (Invitrogen) was used to generate cDNA from 2.0 μg of RNA. Real-time quantitative PCR reactions (15 μl each) contained 1x LightCycler 480 Probes Master Mix (Roche), 2.25 pmol of primers (see Supplemental Table 1 for sequences), and 2 µl of a 1:10 cDNA dilution (1:1000 dilution for GAPDH and mCh). The real time PCR program was run as follows: Pre-incubation, ramp at 4.4°C*sec^−1^ to 95°C, hold 10 min.; Amplification, 45 cycles (ramp at 4.4°C*sec^−1^ to 95°C, hold 10 sec., ramp at 2.2°C*sec^−1^ to 60°C, hold 30 sec., single acquisition); Cooling, ramp at 2.2°C*sec^−1^ to 40°C, hold 30 sec. Crossing point (C_p_) values, the first peak of the second derivative of fluorescence over cycle number, were calculated by the Roche LightCycler 480 software. Expression level was calculated as *delta C*_*p*_ = 2^[*C_p_ GAPDH* - *C*_*p*_ *experimental gene*]. Fold change was determined as *double delta C*_*p*_ = *delta C*_*p*_ *treated cells* / *delta C*_*p*_ *mock* for PcTF expression levels (Fig. 3C), or as *double delta C*_*p*_ = *C*_*p*_ *dox treated cells* / *delta C*_*p*_ *no dox* for gene expression levels in the stable cell lines (Fig. 3D).

### Transcriptome profiling with RNA-seq

RNA-seq was performed using two biological replicates per cell type, treatment, and time point for transiently transfected cells and three replicates for untransfected MCF10A. Total RNA was prepared as described for qRT-PCR. 50 ng of total RNA was used to prepare cDNA via single primer isothermal amplification using the Ovation RNA-Seq System (Nugen 7102-A01) and automated on the Apollo 324 liquid handler (Wafergen). cDNA was sheared to approximately 300 bp fragments using the Covaris M220 ultrasonicator. Libraries were generated using Kapa Biosystem’s library preparation kit (KK8201). In separate reactions, fragments from each replicate sample were end-repaired, A-tailed, and ligated to index and adapter fragments (Bioo, 520999). The adapter-ligated molecules were cleaned using AMPure beads (Agencourt Bioscience/Beckman Coulter, A63883), and amplified with Kapa’s HIFI enzyme. The library was analyzed on an Agilent Bioanalyzer, and quantified by qPCR (KAPA Library Quantification Kit, KK4835) before multiplex pooling and sequencing on a Hiseq 2000 platform (Illumina) at the ASU CLAS Genomics Core facility. Samples were sequenced at 8 per lane to generate an average of 2.5E+07 reads per sample. Read values ranged from 5.7E+06 (minimum) to 1.11E+08 (maximum) per sample.

### Transcriptome analysis

RNA-seq reads were quality-checked before and after trimming and filtering using FastQC [112]. TrimmomaticSE was used to clip bases that were below the PHRED-scaled threshold quality of 10 at the 5’ end and 25 at the trailing 3’ end of each read for all samples [113]. A sliding window of 4 bases was used to clip reads when the average quality per base dropped below 30. Reads of less than 50 bp were removed. A combined reference genome index and dictionary for GRCH38.p7 (1-22, X, MT, and non-chromosomal sequences) [114] that included the full coding region of the synthetic PcTF protein were created using Spliced Transcripts Alignment to Reference (STARv2.5.2b) [115] and the picard tools (version 1.1.19) [116]. Trimmed RNA-seq reads were mapped, and splice junctions extracted, using STARv2.5.2b read aligner [115]. Bamtools2.4.0 [117] was used to check alignment quality using the ‘stats’ command. Mapped reads in BAM format were sorted, duplicates were marked, read groups were added, and the files were indexed using the Bamtools 2.4.0 package. CuffDiff, a program in the Cufflinks package [68], was used to identify genes and transcripts that expressed significant changes in pairwise comparisons between conditions. Fastq and differential expression analysis files are available at the National Center for Biotechnology Information (NCBI) Gene Expression Omnibus (GEO) database (Accession GSE103520, release date September 8, 2017). CummeRbund [68] was used to calculate distances between features and to generate graphs and charts (JSD plots). R ggplot2 [114,118] and VennDiagrams [119] were used to generate heat maps and Venn diagrams respectively. The entire workflow is provided as a readme file at: https://github.com/WilsonSayresLab/PcTF_differential_expression

### Bioinformatics analyses and sources of publicly shared data

*Chromatin immunoprecipitation followed by deep sequencing (ChIP-seq) data*: For the results shown in Figure 1B, H3K27me3 data for MCF7 cells was downloaded from the ENCODE project (accession UCSC-ENCODE-hg19:wgEncodeEH002922) [120]. We classified genes with a ChIP-seq peak within 5000 bp up or downstream of the transcription start site as H3K27me3-positive (1,146 protein-coding transcripts). EZH2-enriched genes (2,397 protein-coding transcripts) for MDA-MB-231 [16] were provided as a list from E. Benevolenskaya (unpublished). For the results shown in Figure 5 and S6, MCF7 ChIP-seq data (from the P. Farnham, J. Stamatoyannopoulos, and V. Iyer labs) was downloaded from the ENCODE project [120]: H3K27me3 (ENCFF081UQC.bigWig), H3K9me3 (ENCFF754TEC.bigWig), H3K27ac (ENCFF986ZEW.bigWig), H3K4me3 (ENCFF530LJW.bigWig), and RNA PolII (ENCFF690CUE.bam) and used to generate plots using DeepTools [121] (computeMatrix, plotProfile, plotHeatmap) in the Galaxy online platform at usegalaxy.org [122]. Prior to plotting, the RNA PolII data was converted to bigWig format using bamCoverage. *Gene ontology term enrichment*: GOrilla analysis used the following parameters: organism, *Homo sapiens*; mode, target and background ranked list of genes; ontology, process; *p*-value threshold = 10.0E-3) [72]. The background ranked list is available at https://github.com/WilsonSayresLab/PcTF_differential_expression. Panther analysis used the following parameters: analysis type, PANTHER Overrepresentation Test (Released 20171205); annotation version, PANTHER version 13.1 Released 2018-02-03; reference List, *Homo sapiens* (all genes in database); annotation data set, PANTHER GO-Slim biological process. Figure 3C was generated using REViGO [123] and GOrilla. Unique differentially expressed genes were analyzed using GeneCards [93]*. Promoter motif analysis*: The script TF_targets was downloaded from https://github.com/cplaisier/TF_targets and used to find enriched transcription factor target sites that were determined by empirical evidence from chromatin studies across 68 cell lines[76]. *Tumor suppressor genes*: The results in Figure 7 are based on human tumor suppressor genes (983 total) that are reported to show lower expressed in cancer samples of the Cancer Genome Atlas (TCGA) compared to the TCGA normal tissue samples was downloaded from https://bioinfo.uth.edu/TSGene/download.cgi. Of these 983 genes, 589 are breast cancer specific [91,92].

## ACKNOWLEDGEMENTS

The project was supported by a grant from the Arizona Department of Health Services (ADHS) Arizona Biomedical Research Commission (ABRC) (14-082976 to K.A.H.). M.A.W.S. and K.C.O. were supported by startup to M.A.W.S from the School of Life Sciences and the Biodesign Institute at Arizona State University. K.A.H. was supported by K01 CA188164 from the NIH NCI. D.B.N. was supported by 14-082976 from the ADHS. The authors thank E.V. Benevolenskaya for the gift of the MDA-MB-231 EZH2 gene module list and C. Plaisier for help with TF_targets.

## AUTHOR INFORMATION

### Contributions

K.C.O. performed differential expression transcriptome analysis, identification of targeted upregulated genes in response to PcTF, and submission of NGS data to Gene Expression Omnibus (GEO). D.B.N performed cell culturing and transfection, preparation of samples for RNA-seq, and RT-qPCR. D.A.V. completed transcription factor motif analyses. M.A.W.S. was responsible for the oversight of the bioinformatics analyses and interpretation of the data. K.A.H. was responsible for the conception of the project, and oversight of molecular cloning, cell culturing, and RNA-seq. All authors contributed to writing of the manuscript.

